# A multi-reporter bacterial 2-hybrid assay for the fast, simple, and dynamic assay of PDZ domain – peptide interactions

**DOI:** 10.1101/512566

**Authors:** David M. Ichikawa, Carles Corbi-Verge, Michael J. Shen, Jamie Snider, Victoria Wong, Igor Stagljar, Philip M. Kim, Marcus B. Noyes

**Author notes:** These authors contributed equally (David M. Ichikawa, Carles Corbi-Verge).

## Abstract

The accurate determination of protein-protein interactions is an important goal of molecular biology and much progress has been made with modern methods emerging in the past decade. However, current methods have limitations, including scale and restriction to high affinity interactions. This has limited our understanding of the large subset of protein-motif interactions. Here we describe a modified bacterial-hybrid assay that employs a combined selectable and scalable reporter system that allows the screen of large protein-peptide libraries and their sort by relative strength. We have applied this tool to characterize a set of human and *E. coli* PDZ domains. Our results are consistent with prior characterization of these proteins and the improved sensitivity increases our ability to predict known and novel *in vivo* binding partners. This approach allows for the recovery of a wide range of affinities with a high throughput method that does not sacrifice the scale of the screen.

---

Protein interactions are at the foundation of cellular processes yet our understanding of how variation across the proteome may influence these interactions is cursory at best. In fact, it is estimated that there are over 500,000 protein-protein interactions (PPIs) predicted to occur in a human cell^1^ yet we only understand the mechanism behind a very small fraction of these interactions. Understanding these mechanisms may help us to predict the functional consequence of protein variations in these PPIs, a critical point to comprehend as we now know from studies such as the 1000’s Genomes Project that variation is prevalent from one human genome to the next^2,3^. These variations might alter the coding sequence of interacting partners^4^ as well as influence expression levels through modification of regulatory sequences. While we already know that common PPIs play important roles in cellular programs^5^, and dysfunctional interactions are at the heart of many diseases^6^, our limited understanding of the mechanisms behind most PPIs severely restricts our ability to understand how protein variants might influence these interactions.

Understanding the mechanism of any protein interaction can be difficult *a priori*. However, some of the most common interactions are dictated by conserved domain families that have evolved across Archaea, Bacteria, and Eukaryota. Some of these protein domains mediate interactions with short conserved peptide sequences commonly referred to as short-linear motifs (SLiMs), typically comprised of <12 amino acids in disordered regions^7^. Such interactions are of somewhat simpler nature than those of two globular protein domains and are particularly common in signaling pathways. SLiM interaction domains (SLiDs) are extremely common in metazoan proteomes with over 1000 human proteins containing at least one of the top 5 most common (PDZ, SH3, SH2, WD40, and TPR). While some SLiDs require post-translational modification of their target peptides^8^, many do not. For example, the PDZ domain (named after the 3 proteins for which it was first characterized PSD-95/Dlg/ZO-1) primarily specifies the unaltered C-terminal residues of its binding partner though phosphorylation can block or modify this interaction^9^. Unlike eukaryotic proteomes, SLiDs appear far less common in bacteria with the minor exception of the PDZ domain that is found in many prokaryotic species^10^. However, prokaryotes that do express proteins with PDZ domains typically have a small number of these proteins or none at all. Nevertheless, the sheer number of prokaryotic species leads to an extremely large total number of prokaryotic PDZs^11^. In fact, there are more PDZ domains found in 96 common microbes of the human microbiome^12^ (368) than human PDZ domains (266^13^). The prevalence of the PDZ domain in humans and its simplified interaction with the C-terminus, has led to several focused studies^14–17^ but many of these works have been biased by affinity, limited in scale, and largely overlook the prokaryotic PDZs.

Many high throughput screens have been employed to define PPIs^18–24^, however, genetic screens and hybrid assays, in particular driven in a high-throughput manner, have had relatively low sensitivity in detecting motif-mediated interactions. Likewise, affinity-purification mass spectrometry (AP/MS) is focused by design on high-affinity interaction and detects few SLiD interactions. Other methods, are geared directly towards domain-motif interactions, but suffer from other shortcomings. For instance, peptide arrays^23^ are limited by scale, as it’s still relatively expensive to synthesize the peptides on the array, only a limited number of peptides can be measured. Similarly, the hold-up assay offers quantitative affinity data, but is limited by the number of peptides measured^25^. Peptide phage display is another commonly employed method naturally suited to characterize domain-motif interactions^14, 15, 26^. However, as a method based on enrichment over multiple rounds, the results can be biased by affinity and specific amino-acid properties^27^ limiting their predictive power of lower affinity interactions that might be common *in vivo*. In addition, as purifying each individual protein to screen can be laborious, any method that requires protein expression/purification would be quite challenging to employ for an exhaustive screen of PPIs and their variants. Moving forward, we will need a simple, fast and accurate method to address protein-protein variation and provide a predictive model of protein function.

Here we describe a simple, sensitive, multi-reporter bacterial 2-hybrid (MR-B2H) system capable of capturing a dynamic range of protein interactions and enabling the accurate prediction of known and novel *in vivo* protein partners. Using libraries that represent both naturally occurring as well as purely random peptides, we characterize PDZ-peptide interactions that represent all the C-termini a PDZ domain will encounter in a cell as well as all mutant variants of those termini. We apply the MR-B2H system to characterize several PDZ domains from human and *E. coli* proteins that have been investigated by other common methods allowing for the benchmark of our results. We find the MR-B2H results demonstrate an improved ability to predict known *in vivo* interacting partners. The MR-B2H is a robust, rapid, and accurate method that could be adopted by almost any lab. Some of the main benefits of the MR-B2H detailed here are (1) facile workflow circumventing the need for protein purification and using only basic molecular biology reagents (2) parallel and rapid characterization of protein domains, easily screening >15 domains in two-three days (3) accurate prediction of protein interactions (4) sensitive fluorescent output captures a broad range of PPI affinities and allows an approximation of relative activity. While we have applied this technique towards the PDZ domain it could be easily modified to investigate other SLiDs as well as more complex protein interactions.

## RESULTS AND DISCUSSION

### Adapting the multi-reporter bacterial 1-hybrid (B1H) to a B2H

The omega (ω)-based B1H is a well-established, robust, and rapid method for characterizing protein-DNA interactions^28–30^. The strength of this system is that assays can be done in a matter of days and multiple domains can be investigated in parallel. In addition, the system employs the ω-subunit (rpoZ) of bacterial RNA-polymerase (RNAP) as the activation domain and assays are conducted in an *E. coli* strain lacking the endogenous omega as this RNAP subunit is non-essential. In this way recruitment of the polymerase takes place in the absence of competition from the endogenous omega^30^ resulting in a highly sensitive assay of the protein-DNA interaction. In order to adapt this system to characterize PPIs and take advantage of the sensitivity of the omega recruitment we made the following modifications. First, the PDZ domain of interest is expressed as a fusion to the omega subunit from the low copy reporter plasmid (**Figure 1a**). Second, a 7-amino acid peptide is expressed on the C-terminus of the Zif268 DNA-binding domain. The consensus Zif268 binding-site is placed 21 base pairs upstream of a weak minimal promoter that drives the tandem HIS3 and GFP of the reporter plasmid. In this way, activation of the B2H system and expression of the selectable markers HIS3 and GFP is dependent on the PDZ interacting with the peptide: The DNA-binding domain interacts with its target and omega will interact with the RNAP, but the two components will only be brought together on the reporter plasmid through a positive PDZ-peptide interaction. Further, the strength of the PPI required to survive selective pressure can be titrated by different concentrations of 3-amino triazole (3AT) – a competitive inhibitor of the HIS3 gene product.

**Figure 1.**
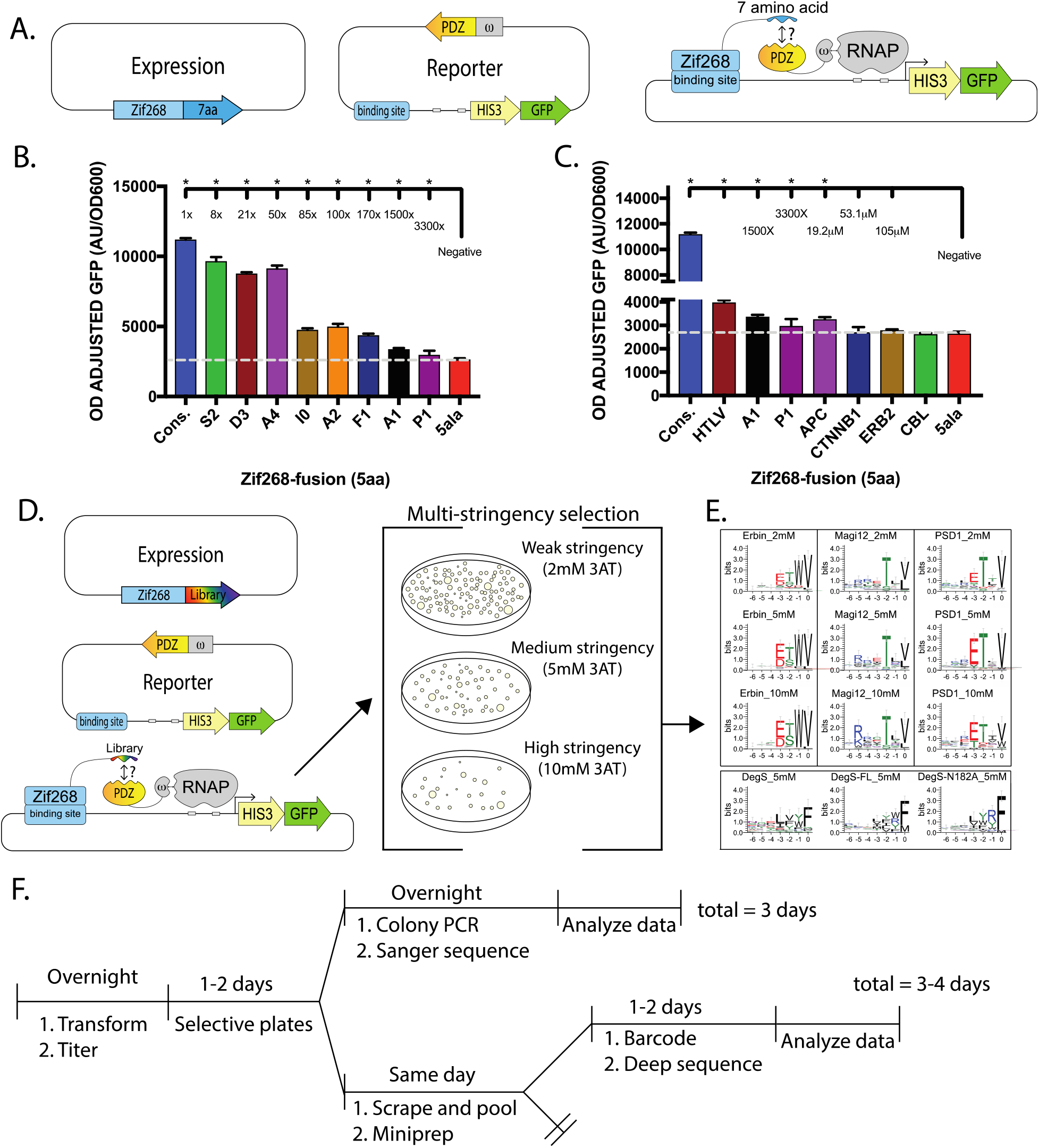
A multi-reporter bacterial 2-hybrid (MR-B2H) tool for the selection of PDZ-peptide interactions. **A)** Vectors used and selection scheme are shown. A positive interaction between a Zif268-peptide and an omega-PDZ domain recruits RNA-polymerase (RNAP) and drives activation of the reporters HIS3 and GFP. **B)** GFP expression driven by known ERBIN-peptide interactions. The reduction in IC50 measurements are shown at the top for each peptide. The peptide labels represent the amino acid substituted at which position of the consensus target. **C)** GFP expression driven by the C-termini of proteins the ERBIN PDZ is known to interact with. Affinities of some known interactions are noted above. **D)** Scheme of a selection of positive PDZ-peptide interactions on plate with varying concentrations of 3-AT. **E)** Motifs of pooled surviving colonies from plate selections at 2, 5, and 10mM 3AT. Bottom: Selections of prokaryotic PDZ domains and full-length proteins. **F)** Timeline of workflow.

To test the functionality and demonstrate the sensitivity and dynamic range of the MR-B2H, we tested nine previously characterized peptides paired with the PDZ domain of the ERBIN protein for their combined ability to activate our GFP reporter. These peptides were chosen from prior work as they had demonstrated a wide-range of IC50 values (3,300 fold)^31^ when bound to the ERBIN PDZ. Cells that expressed the omega-Erbin PDZ domain and the zif268-peptide fusion were challenged for their ability to activate the GFP reporter. Known lower-affinity interactions drove less GFP expression than high-affinity interactions (**Figure 1b**), but were still distinguishable from negative controls. Importantly, documented ERBIN PDZ-peptide interactions found in the mentha database^32^ clustered in the lowest tier of GFP expression (**Figure 1c**). This data suggested that the MR-B2H system captures a wide range of affinities, and that many relevant interactions would be found in the lower GFP range.

### Characterization of PDZ specificity

Having demonstrated the ability to capture a wide range of affinities, we next tested our ability to select positive PDZ-peptide interactions from large libraries. We constructed a purely <u>random</u> 7 amino acid library as well as a library comprised of ~50,000 <u>naturally</u> occurring 7 amino acid C-termini found in humans (templated from prior work)^15^. Combined, these libraries allow us to investigate peptides a PDZ would be exposed to in nature in addition to all possible variations of these peptides. While the build of the random library approached 10^10^ variants, the 10^9^ transformation efficiency of *E. coli* limits the practicality of sampling fractions of this library greater than 10^9^ in a single experiment. Therefore, all selections reported here are based on a sampling of roughly 10^9^ members of our random library. The natural library of only 50,000 members was easily oversampled with a typical sampling 5×10^7^ variants in each selection, representing a 1,000-fold over sample.

To demonstrate the method’s ability to select positive interactions from our libraries and generate results rapidly, we carried out MR-B2H selections as previously described for B1H selections^30,33^ relying only on HIS3 activation. Selections were plated on a range of 3-amino triazole (3AT) concentrations, and individual colonies were screened for surviving library members by colony PCR or pooled and barcoded for Illumina sequencing (**Figure 1d**). As might be expected, the pools of library members recovered from lower stringency selections were more diverse than those from higher stringency selections. We initially performed selections against 4 PDZ domains from eukaryotic proteins, and 6 PDZ domains from 4 prokaryotic proteins. 3/4 eukaryotic PDZ domains yielded specificities matching known binding preferences (with Scribble PDZ 3 having too few colonies to obtain reliable information), and at least one PDZ worked for 3/4 prokaryotic proteins (with the PRC PDZ domain failing selection) (**Figure 1e**, and **Supplemental Figure 1**). Importantly, the motif from DegS PDZ alone matches closely to the motifs^34^ from the full-length protein (expressed as a catalytically active wild-type, or catalytically impaired mutant^35^) indicating that a domain-based screening approach can be informative of full-length protein binding as well. This entire workflow can be accomplished in as little as 3 days (**Figure 1f**), highlighting this method as a rapid and facile approach for characterizing SLiDs.

### The B2H system captures a wide range of affinities

To expand on the multi-stringency selections, we took a complementary approach to leverage GFP expression as a relative measure of interaction strength. Rather than relying on 3AT stringency to select strength of interaction in a step-wise manner, we used GFP fluorescence as a continuous measure of interaction strength. However, to clear nonfunctional members from the library we first needed to perform a weak 3AT selection on plates (typically 1 mM 3AT). We then pooled surviving members and expanded in liquid media under the same weak selective pressure until GFP fluorescence approached its maximum. The GFP populations were sorted representing low, medium and high GFP (**Figure 2a**). A negative GFP population was also recovered to provide a baseline control. Consistent with the results from the plate selections, we found a more diverse set of peptides in lower GFP populations, while a more restricted set of peptides were required for maximal GFP activation (**Figure 2b**). In fact, we find that in general, motifs compiled from lower GFP bins more closely resemble the diversity of interacting partners that are found *in vivo*. For example, the low GFP results for the ERBIN PDZ domain more closely resemble results from peptide array^36^ as well as confirmed *in vivo* binders curated from *mentha^32^* (**Figure 2b bottom**). On the other hand, motifs from the high-GFP expressing populations lead more towards the high affinity consensus targets produced by phage-display. Further support of these trends relating affinity and GFP comes from the distribution of peptides with documented affinities across GFP. Consistent with results above, we find an enrichment of lower affinity interactions (>50uM) in low GFP pools, and higher affinity interactions (<10uM) in the high GFP populations (**Figure 2c**). Utilizing the GFP component of the MR-B2H enhances the data by providing a continuous separation of PPI strengths but adds an additional 4-5 days to the workflow described in **Figure 1f**, (**Figure 2d**).

**Figure 2.**
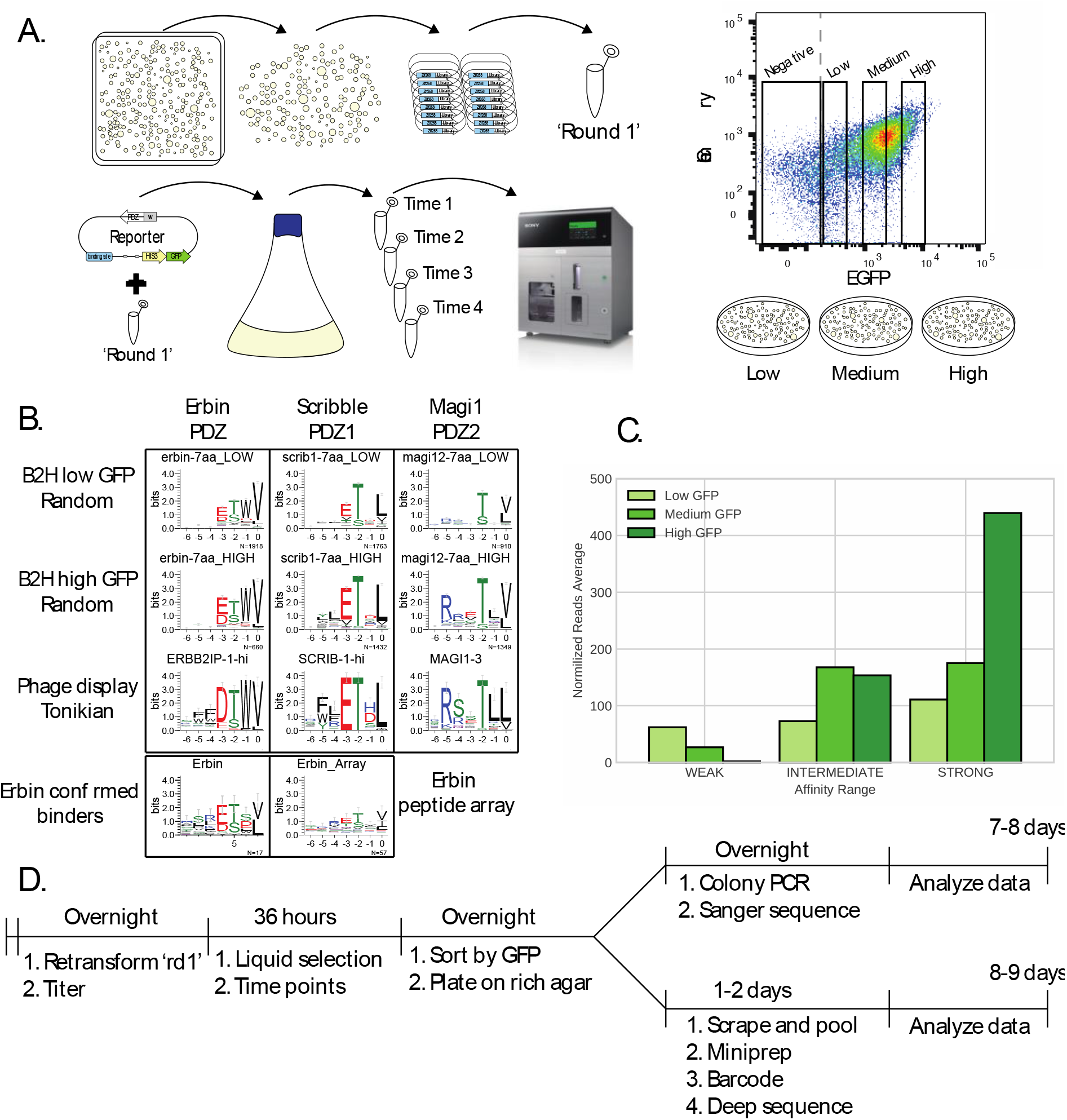
The MR-B2H system captures a wide range of functional affinities. **A)** Transformations are performed as in **Figure 1d**. Surviving colonies were and pooled and miniprepped to obtain an enriched library termed ‘Round1’. Retransformed DNA ies grown in liquid media until GFP approaches maximum fluorescence (typically 36 hours). Three populations of cells with approximately 5-fold differences in mean GFP fluorescence are sorted and sequenced. **B)** Motifs recovered in low and high GFP populations are shown and compared to phage display result and known *in vivo* binding partners. **C)** Twenty-five proteins with known affinities for either the ERBIN or the Scribble PDZ domains were identified by the MR-B2H screen and the distribution of the reads for these peptides across GFP populations is plotted. Weak affinities are >50uM, mild affinities = 10-50uM, and strong affinities are <10uM. **D)** A simplified timeline for experiment is depicted. This workflow picks up where **Figure 1f** ends.

This workflow was performed for 9 previously characterized PDZs from 6 eukaryotic proteins, including the ERBIN PDZ, DLG4 PDZ3, SCRIB PDZ1 and PDZ3, MAGI1 PDZ2, 3, and 4, MPDZ PDZ10 and TIAM1 PDZ1. We successfully measured the specificities of 8/9 of these PDZs in good agreement with previous work, with the TIAM1 PDZ1 being the only PDZ that failed in the B2H assay. Additionally, 3 PDZs which had been attempted in phage display but failed to produce motifs were successfully characterized, including DLG4 PDZ1 and PDZ2, and APBA1 PDZ1 (**Supplemental Figure 2 and 3**).

### Predictions of known interactions are improved by models incorporating lower affinity binders

One goal of this project is to produce a system that captures a more representative group of peptides that interact with domains like PDZ’s in order to improve our capacity to predict novel interactions. Utilizing published phage display results as a comparison^15^, we interrogated the data from our selections of the Natural Library. After filtering selection data to enrich for high confidence binders (see Methods), we plotted the distribution of recovered known “annotated” binders across GFP populations (**Figure 3a**). Annotated binders are found in all three GFP populations, however in most cases, the lower GFP populations contain a greater proportion of these peptides, supporting the notion that there is a wealth of lower-affinity interactions that may be overlooked by methods biased towards the high affinity. Furthermore, the absolute recovery of known interactors by MR-B2H selections is 4-10-fold higher than phage display (**Figure 3b**). To control for false discovery rates and directly compare to phage display results, we then built position weight matrices (PWMs) from MR-B2H-derived peptides found in either low or high GFP populations, as well as phage display-derived peptides, and scored each C-terminus of known partners. With a false-discovery rate set to 0.01, we found that in most cases, the highest number of known binders passed the cutoff when compared to the PWM derived from low-GFP B2H peptides (**Figure 3c**). Stated simply, PWMs derived from a broader set of PPI affinities and characteristics recover a higher number of known binders than PWMs based on more restrictive information. Phage-display-derived peptides are known to be strong and hydrophobic binders^27^, but lack the affinity range captured by MR-B2H selection, hence binding profiles (e.g., expressed as PWMs) derived from them often fail to capture important interactions. In sum, these results demonstrate that incorporating a richer spectrum of interaction affinities into our models significantly improves our ability to predict known interactions.

**Figure 3.**
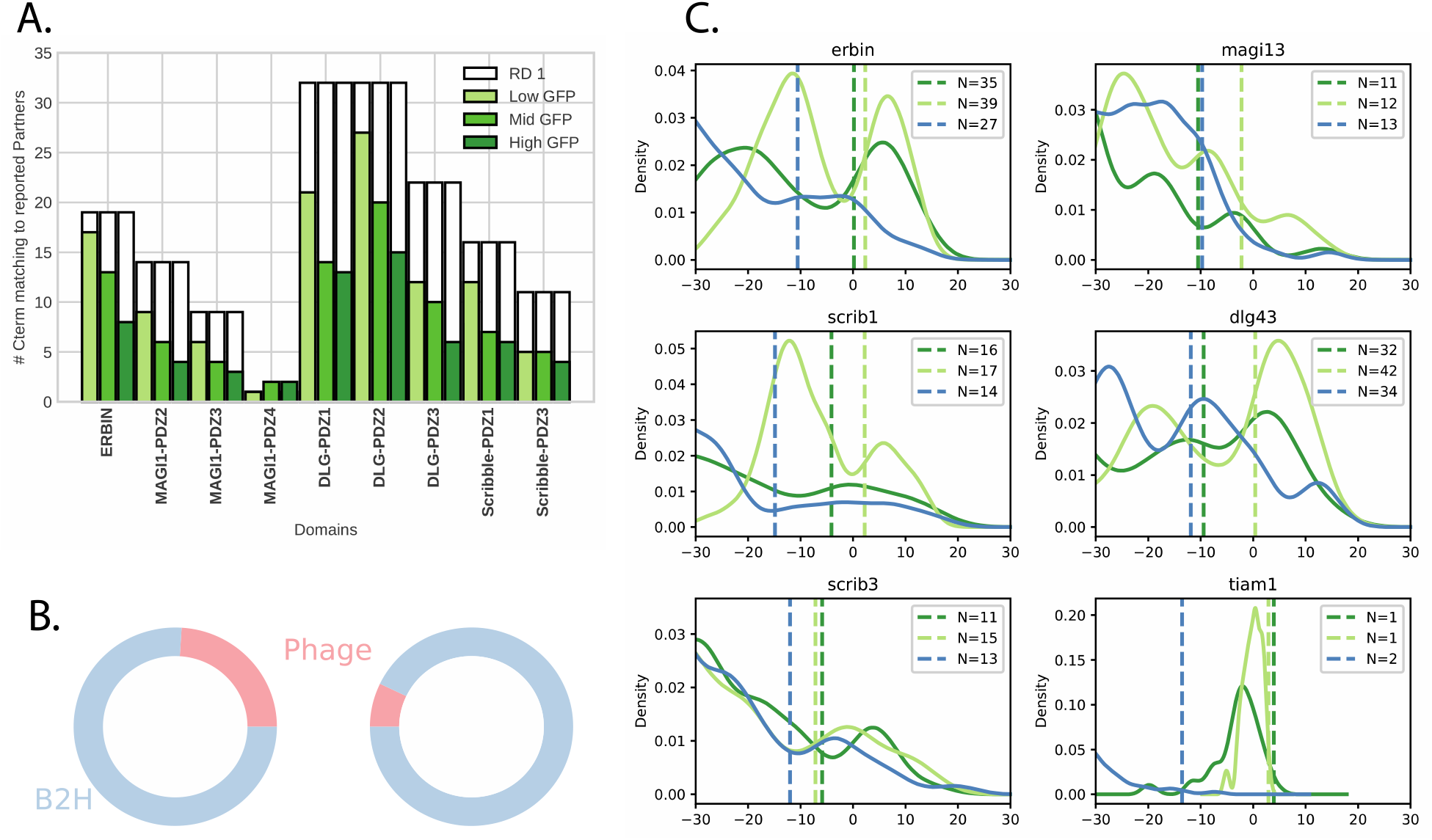
Capture of broad GFP outputs improve the accuracy of predicting known PDZ interacting partners. **A)** Plotted counts of selected peptides matching C-termini of annotated *in vivo* partners. Empty box indicates total annotated C-termini passing the reads-threshold after round 1 selection. Green bars depict the number of annotated matches within each GFP level. **B)** Comparison of the absolute number of matches to previously reported PDZ partners recovered by either Phage Display or MR-B2H selection. **C)** Comparison of the number of C-termini identified as binders using a PWM built using phage-display selected peptides (blue line) and MR-B2H-selected peptides from the Random library for low and high GFP (light and dark green lines, respectively). The C-termini of all previously described partners was scored. The threshold for each PWM was set to ensure an FDR of 0.01 for phage display and MR-B2H PWMs (dashed lines).

### The prediction of novel interaction partners underscores the utility of MR-B2H data

The agreement between our MR-B2H data and previous screens carried out by phage display^14,15^ and protein microarrays^16,36^ validates the accuracy of our technique. In addition, the improved prediction of known *in vivo* binding partners with MR-B2H data (**Figure 3**) demonstrates the importance of the expanded affinity profile for detecting real full-length protein interactions. However, many recovered peptides were not from known interacting partners and we next set out to determine if these represent false positive or novel true positive interactions. That said, it is important to note that the possibility of interaction does not imply biological function or purpose but simply that if two proteins are present in the same cell they have the potential to interact. Further, it is beyond the scope of this work to thoroughly investigate the biological implications of each novel prediction of PDZ and peptide pairings. Rather, we aim to demonstrate this tool is able to accurately predict the possibility of *in vivo* interacting partners that others might follow up in future works. With this goal in mind, we turned to another hybrid assay that would express a set of full-length proteins in human cells to test for positive interactions.

Since the PDZ domain is often found in proteins orchestrating scaffolding interactions at the cell membrane^37^, we chose to focus our human cell validation on the PSD-95 (gene name DLG4), a neuronal membrane protein containing 3 PDZ domains. We characterized the target specificity of each PSD-95 PDZ domain (**Supplemental Figures 2 and 3**) and not surprisingly found that many of the predicted partners map to intermembrane proteins. To validate these predictions, we used the mammalian membrane 2-hybrid system (MaMTH)^38–40^ to investigate a small set novel predicted partners (**Figure 4a**). This system utilizes the reconstitution of a split ubiquitin molecule that only happens when two interacting partners brings these halves in close proximity to one another. The reconstitution leads to cleavage of a transcription factor and its subsequent transport to the nucleus where it activates the reporter gene luciferase. In this way, luciferase activation is a measure of the interacting partners at the membrane. Using this method, we tested a set of five predicted partners (four novel: ACVR2B, LRNF4, ATP2B2, and SLC2A1. One known: FZD7) with each of the PDZ domains alone as well as the full-length protein (**Supplemental Figure 4**). Luciferase activity was significantly increased over the controls for all of these interactions with the full-length protein (**Figure 4b**) and for 4/5 interacting partners with the single PDZ domains (**Supplemental Figure 4**). Interestingly, while still significantly above controls, the luciferase activity decreased for 4/5 of the interactions with the full-length protein in comparison to any of the individual PDZ domains. In addition, the one exception, SLC2A1, significantly increases luciferase output with the full-length but not a fragment that contains all 3 PDZ domains. These results imply that the PDZ interaction may be very weak as luciferase activity is slightly increased (though not significant) for each PDZ domain, but the more than doubling of reporter output with the full-length protein is likely due to an additional interaction in the protein. This secondary interaction may or may not require the weak PDZ interaction, but determining this interplay would require detailed follow up investigations. Regardless, these results demonstrate that the MR-B2H data was able to successfully predict 4/4 novel full-length interacting partners with PSD-95.

**Figure 4.**
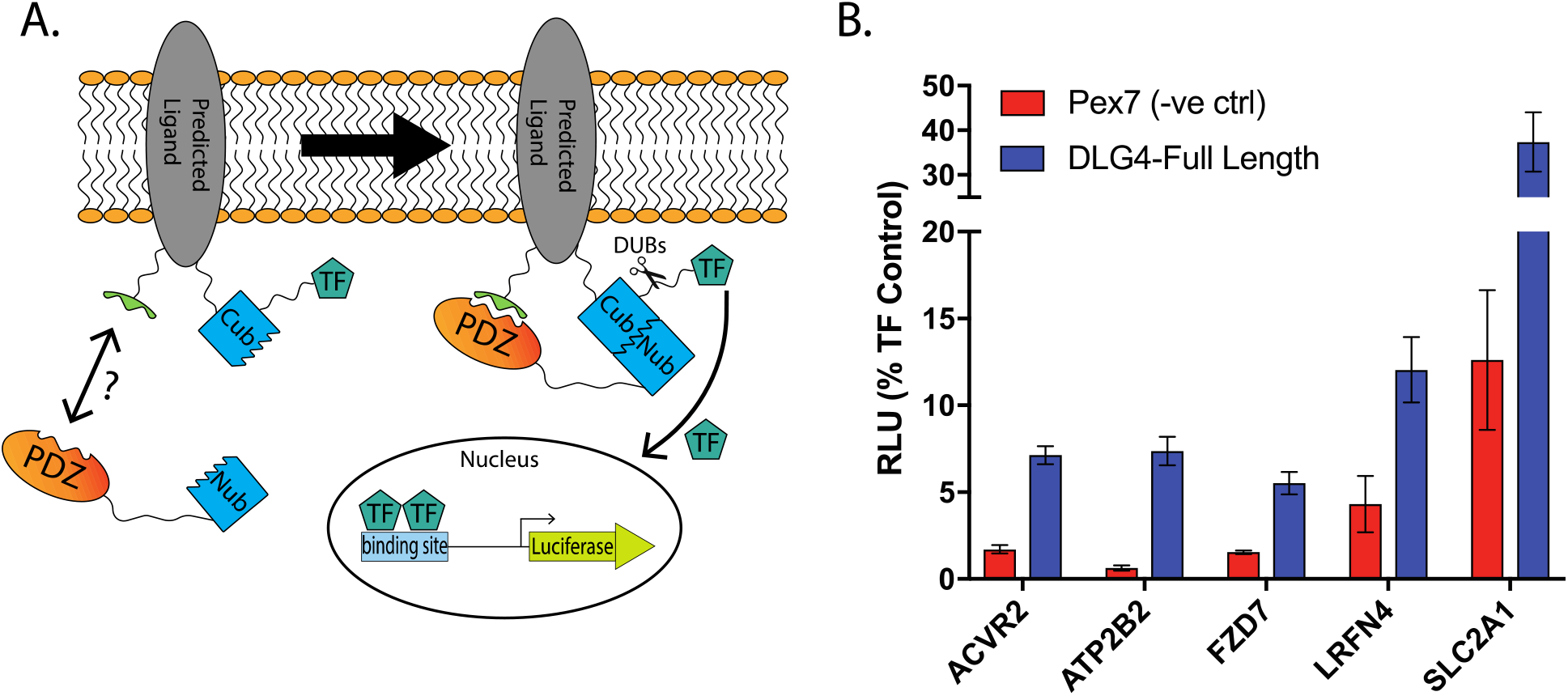
Specificities produced by the MR-B2H system improve the prediction of novel PSD-95 interacting partners at the cell membrane. A) Schematic of the mammalian membrane 2-hybrid (MaMTH). If the PDZ domain interacts with the C-terminus of its partner, a pseudo ubiquitin molecule is reconstituted. De-ubiquitinases cleave off the fused transcription factor, which translocates to the nucleus and activates expression of a luciferase reporter. B) MaMTH luciferase expression as a fraction of a control transcription factor with a known PSD-95 interaction partner (FZD7) and 4 MR-B2H predicted partners.

### The MR-B2H provides insight into the specificities of bacterial PDZ domains

The specificities of prokaryotic PDZ domains have been largely ignored outside of a handful of E. coli proteins. One potential complication for the prokaryotic domains is the genes that code for them often contain a protease domain. Expressing a functional protease in a hybrid assay could prove problematic. However, structures of the DegS proteins reveal that the protease domain is in close proximity to the peptide bound to the PDZ domain^41^ indicating that this interaction may influence specificity. Therefore, to test the ability of the MR-B2H system to characterize these prokaryotic domains we first focused on the DegS protein. Using the Random library we tested DegS as the PDZ domain alone, as the full-length protein including the protease domain, and a mutant version of the protein that reduces the catalytic activity of the protease (**Figure 5**). The DegS peptide preference is in agreement with prior investigation of this domain^34^. Further, we find only subtle differences between the 3 versions of the proteins with the full-length versions enriching for an Arginine at the −1 position not found with the PDZ domain alone. Therefore, the PDZ domain alone appears to provide at least an approximation of the full-length specificity. Also, our experiments demonstrated no signs of toxicity when we overexpress this *E. coli* protease in our *E. coli*-based system indicating that investigation of other PDZ-proteases are not likely to be problematic in our system. Next, we screened the PDZ domains of 3 other E. coli proteins DegP, DegQ, and PRC. Only PRC failed to enrich for specific peptides. DegP and DegQ both contain 2 PDZ domains but only one of each produced a motif (**Figure 5**). The three failed domains may require additional sequences to be functional or even expression as full-length proteins but this would make it difficult to distinguish which PDZ was responsible for binding the peptide. Regardless, for the PDZs successfully characterized, their peptide preferences are in agreement with published work^42,43^ validating the MR-B2H system’s ability to characterize prokaryotic domains in addition to their eukaryotic counterparts.

**Figure 5.**
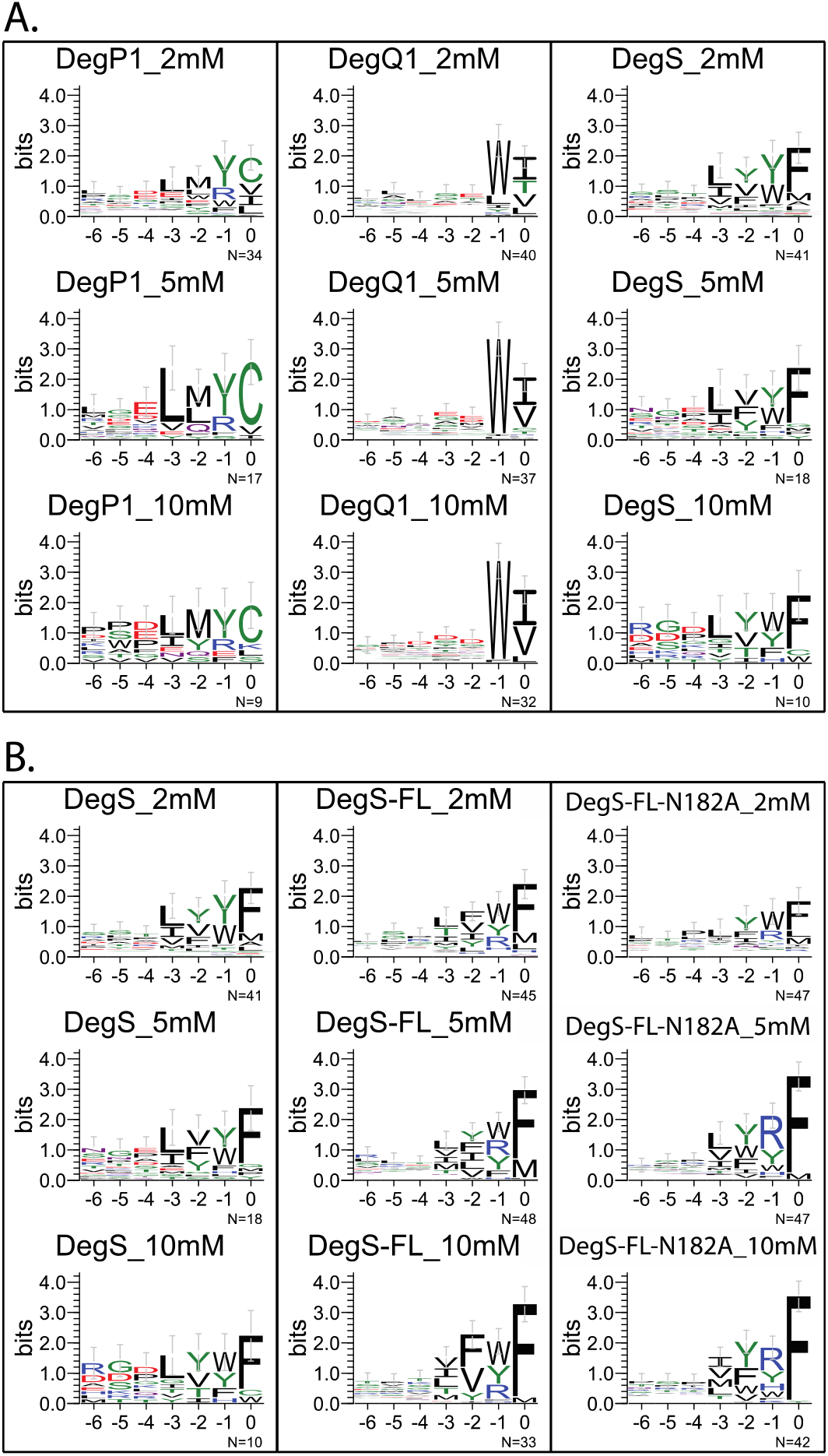
Peptides selected by MR-B2H to bind E. coli PDZ domains. Selections were done on plates at the 3AT stringency indicated above each motif. ***Right***, DegS was tested 3 ways: as the PDZ alone (DegS), the full-length protein (DegS-FL), and a mutant version of the full-length protein (N182A) that significantly reduces the activity of the protease domain.

## Conclusions

Here we describe a rapid, sensitive, and accurate hybrid system enabling the parallel characterization of PPIs in as little as three days. Relative activity of interactions can be obtained in as little as four to five additional days. We showcase the MR-B2H using the PDZ domain, one of the most prevalent SLiDs in human. We successfully apply this method to 8/9 human PDZ domains previously characterized by peptide phage-display, as well as 2 domains that had failed. Additionally, we identify a novel motif for the human APBA PDZ domain. Finally, we characterize 3 PDZ domains from *E. coli* and demonstrate the successful capture of protein-ligand interactions even when these PDZ domains are expressed as full-length proteins and include a functional protease. In all cases the specificities produced are in agreement with prior work (when available) demonstrating the accuracy of the technique. As a result, the combined speed, throughput and accuracy of this method should make it useful for future studies of PDZ domain proteins as well as other common SliDs. In addition, the simplicity of the assay allows for its employment by any lab skilled in basic molecular biology techniques.

The goal of any high throughput technique is to accurately capture the true activity of a protein *in vivo*. Therefore, our goal was to create a tool that would improve our ability to predict *in vivo* binding partners, first for PDZ domains, but with the potential to expand to other protein domains. While prior methods to characterize these domains have sacrificed throughput or sensitivity^14–16, 36^, the MR-B2H system allows the screen of large libraries without sacrificing the capture of a broad range of affinities. We believed that the capture of a wide range of affinities without sacrificing scale would enable better prediction of *in vivo* binding partners. We find this to be true as our MR-B2H data is better able to predict known *in vivo* interacting partners than prior works and our MaMTH studies demonstrate the accurate prediction of novel binding partners.

The application of this method could be beneficial in several ways. First, the simple selection of interactions at high stringency from plates can provide a consensus binding motif for a protein within 3 days. Second, application of low stringency selections, or low GFP recovery, can provide a broader understanding of the domain-peptide interaction. Scaled and applied to a large set of domains, the mechanistic fundamentals of the domain’s function may be gleaned. As compared to phage display, the nature of our method sidesteps the laborious and often difficult need to generate a functional purified protein in vitro. This approach has been applied to many DNA-binding domains^28–30,44,45^ and the sensitivity of the MR-B2H system may enable a similar approach to understand SLiD function. Finally, the simplicity and throughput of this technique allows the seamless application of multiple libraries. The comparison of the natural library with a purely random library allows us to investigate what peptides a protein might interact with in a given cell, as well as how mutations in the peptide might alter the interaction. Therefore, this comprehensive approach allows us to understand how variation in peptides alter protein recognition. Further the GFP assay allows for a rough quantification of how slight variation in the peptide may influence the relative strength of the interaction. As it has become clear that variation across human genomes is prevalent^2^, understanding variation in PPIs, especially those that can influence low affinity interactions, will be critical as we try to decipher whether a variant is harmful or benign.

## MATERIALS AND METHODS

### Library and plasmid construction

The Random and Natural libraries were generated as previously described^46^. Plasmids were generated via routine restriction enzyme cloning, and will be made available at Addgene. Additional details can be found in the Supplemental information.

### Selections on plates

Selections in the B2H assay were performed as previously described for B1H selections^30,33,45^ with cells plated on minimal media (NM) that lacked histidine but supplemented with various levels of 3AT. Surviving colonies were sequenced.

### Selections in liquid culture

To separate positive PDZ-peptide interactions by GFP expression 10^9^ cells were plated on NM media plates lacking histidine but containing 1mM 3AT and grown for 24 hours to enrich for the somewhat rare positive interactions (typically <10^5^ colonies survived out of the 10^9^ plated). Surviving colonies were scraped and DNA was extracted to yield ‘round 1’ libraries. 50ng of round 1 libraries, or the 5 alanine peptide negative control, were retransformed with 1 μg of the same PDZ plasmid. 10^6^ cells were grown in selective NM media supplemented with 1mM 3AT for up to 36 hours. Four GFP populations (negative, low, medium, and high) were sorted and collected from a time-point when GFP approached its maximum. Cells from these sorted pools were plated on rich media and grown overnight. The surviving colonies where pooled, DNA barcoded, and deep-sequenced.

### Next-generation Sequencing

PCR primers that included the illumina adaptor sequences were used to amplify peptide coding sequences from the plate selections as well as recovered GFP populations. Reactions were gel purified and mixed in equimolar ratios before being run a NextSeq4000 with 30% PhiX DNA to increase diversity. Results were de-multiplexed and peptide counts were tallied.

### MaMTH validation assays

MaMTH assays were carried out as previously described^38^. In short, bait and prey plasmids were transfected into HEK293 MaMTH reporter cells, expression was induced with tetracycline, and luciferase activity was measured 24 hours post induction.

### Bioinformatics analysis

For each PDZ domain a PWM was generated from the peptides recovered from each population considered, including all plate stringencies and GFP populations. For comparison, PWMs were also produced from the previously published phage display-derived peptides. In addition, the interacting partners of each target protein were collected from the protein-protein interaction database MENTHA^32^. The C-terminus of each protein annotated as a partner was scored for each PWMs. After scoring, a threshold needed to be set for separating positive and negative predictions. The simplest approach is to set a common log-likelihood threshold for all motifs. Unfortunately, log-likelihood distributions substantially vary across motifs, and the same threshold value may result in underestimating occurrences for one motif and overestimating them for another. Therefore, several score distribution-based approaches for threshold selection were proposed^47^. We selected a method based on controlling the false positive rate, which has proven to be the least biased by the information content of the motif and therefore the most consistent between motifs. Consequently, we compared the performance of each PWM in the dataset of known partners by applying a fixed FDR of 0.01^48^.

The accumulated hydrophobicity value was calculated as the sum of each amino acid hydrophobicity weight, multiplied by each amino acid normalized frequency in the PWM over each position^15^.

## Supporting information

Supplemental Section

## Author Contributions

M.B.N. and P.M.K. conceived of the project. D.M.I. and M.J.S. carried out the bacterial experiments while C.C.V. executed all computational analysis. J.S. and V.W. carried out all MaMTH construct generation and assays. I.S. oversaw all MaMTH assays and provided critical analysis of results. D.M.I., M.B.N. and P.M.K. wrote the manuscript.

## Acknowledgment

NIH T32 #GM066704 supported D.M.I. in this work. PMK and CCV were funded by grants of the Canadian Institute for Health Research (PJT-153279) and the Canadian Cancer Society Research Institute (#702884). MaMTH work performed by the lab of I.S. was supported by CCSRI – 703889, Genome Canada – 9428, Government of Ontario – RE08-9 and ORF-DIG-Stagljar. We thank current and former members of the Noyes lab for the thoughtful suggestions on the work and manuscript. We thank T.M.C. for her valued insight, support and feedback on the manuscript. We would also like to thank the NYU Langone Genome Technology Center for their help in determining the best approach for sequencing our results.

