## Supplemental Section for "A multi-reporter bacterial 2-hybrid assay for the fast, simple, and dynamic assay of PDZ domain – peptide interactions"

1. Department of Biochemistry Molecular Pharmacology and Institute for Systems Genetics, NYU Langone Health, New York, NY 10016, USA.
2. Donnelly Centre for Cellular and Biomolecular Research, University of Toronto, Toronto M5S 3E1, Canada
3. Department of Molecular Genetics, University of Toronto, Ontario, Canada.
4. Department of Biochemistry, University of Toronto, Ontario, Canada.
5. Department of Computer Science, University of Toronto, Ontario, Canada
6. These authors contributed equally
7. Corresponding author

Supplementary Method…………………………………………..Page 8

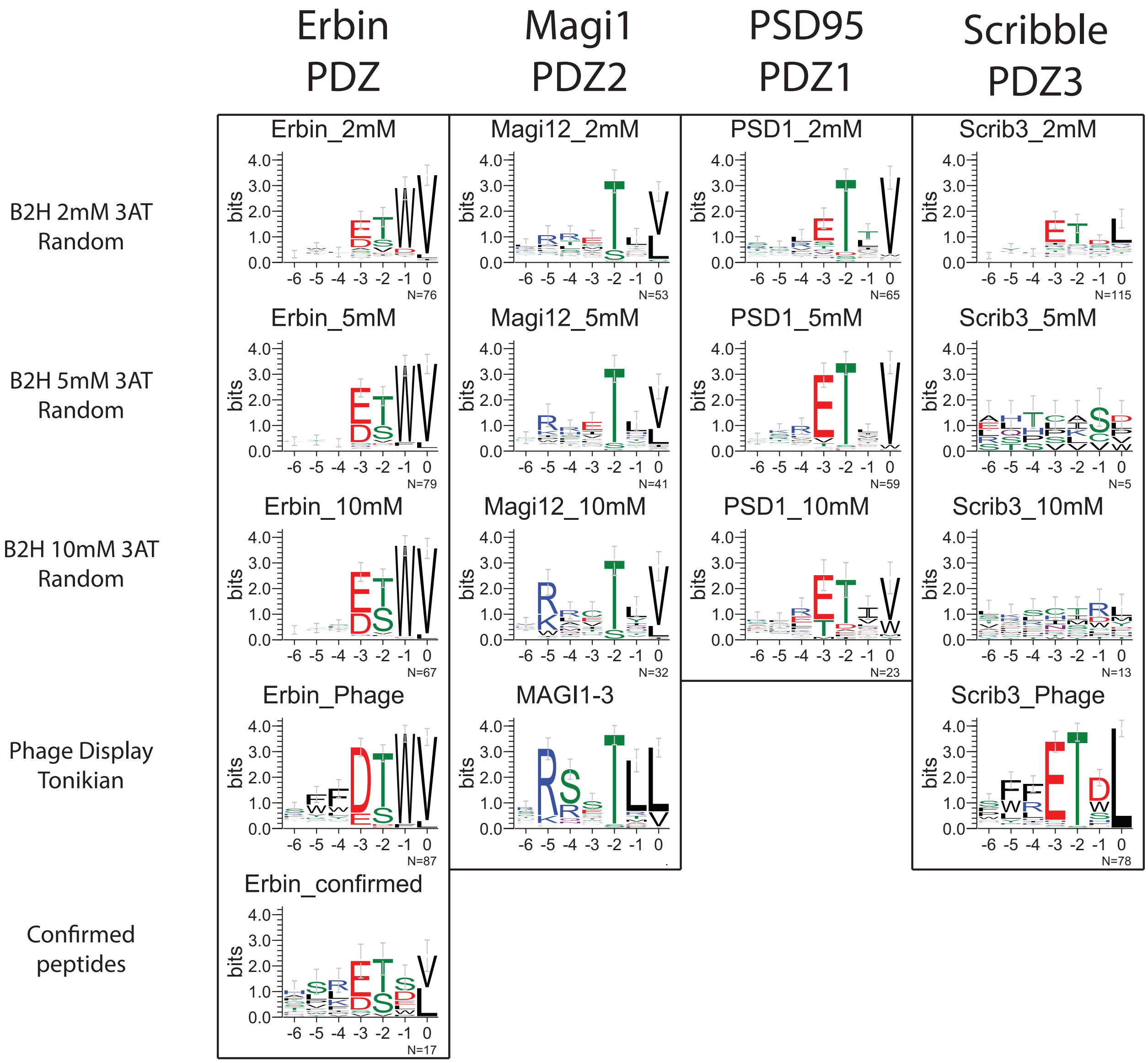

**Figure S1. PDZ specificities determined by MR-B1H <u>plate</u> selections.** *<u>Top 3 rows</u>*, Peptides were selected from the random library by the PDZ domain labeled above each motif. This PDZ domain name is followed by the amount of 3-AT present in the plate for the selection represented. *<u>Bottom rows</u>*, where applicable, motifs are shown that represent peptides selected by phage display, also using a random library[2]. For the Erbin PDZ, a motif of peptides known to bind the domain is also shown[5].

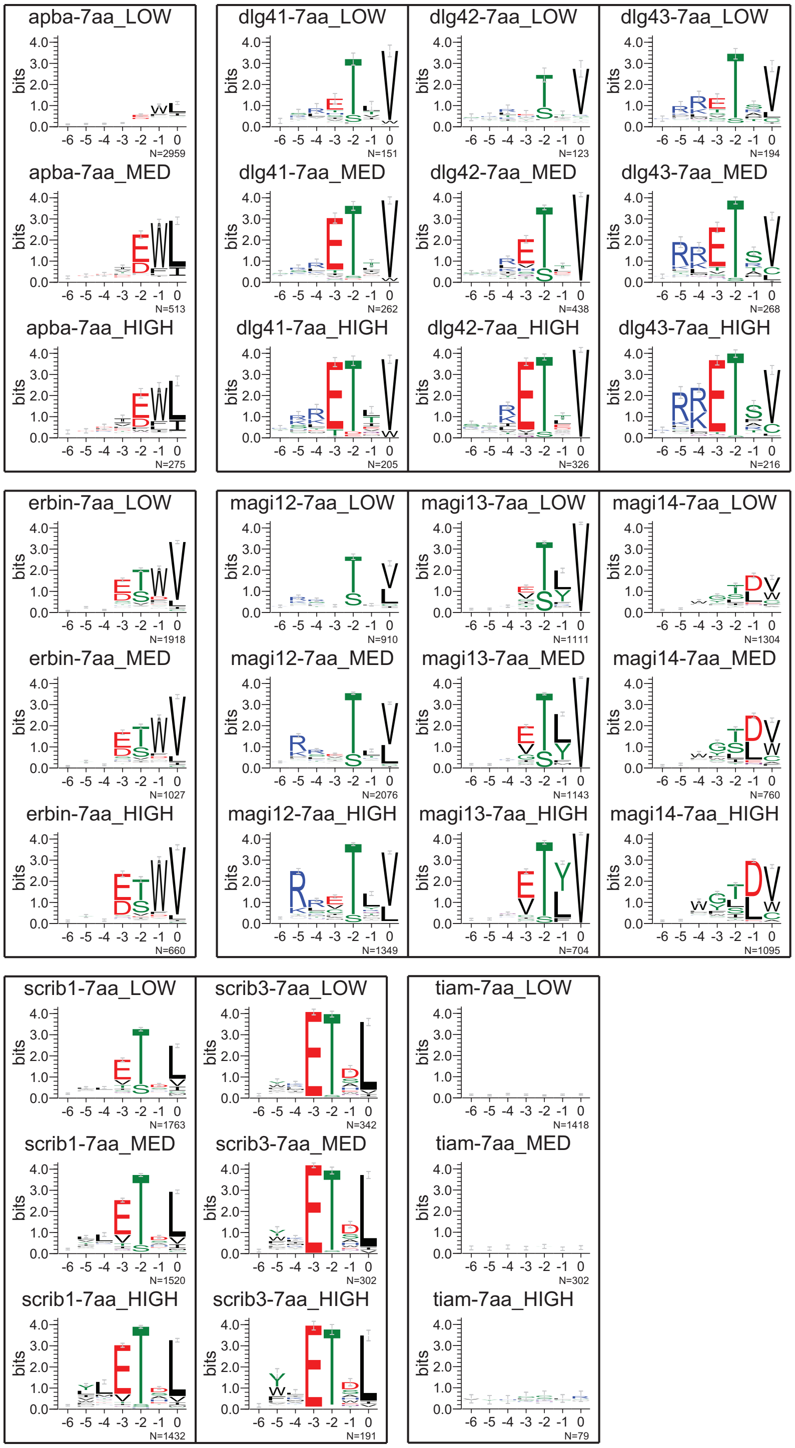

**Figure S2. PDZ specificities determined by MR-B1H selection and <u>GFP sort using a 7aa random library</u>.** Motifs represent peptides selected from the random library and recovered from populations at various GFP levels. The name of the PDZ domain tested is labeled above each motif as well as the GFP level it represents (LOW, MED(medium), or HIGH).

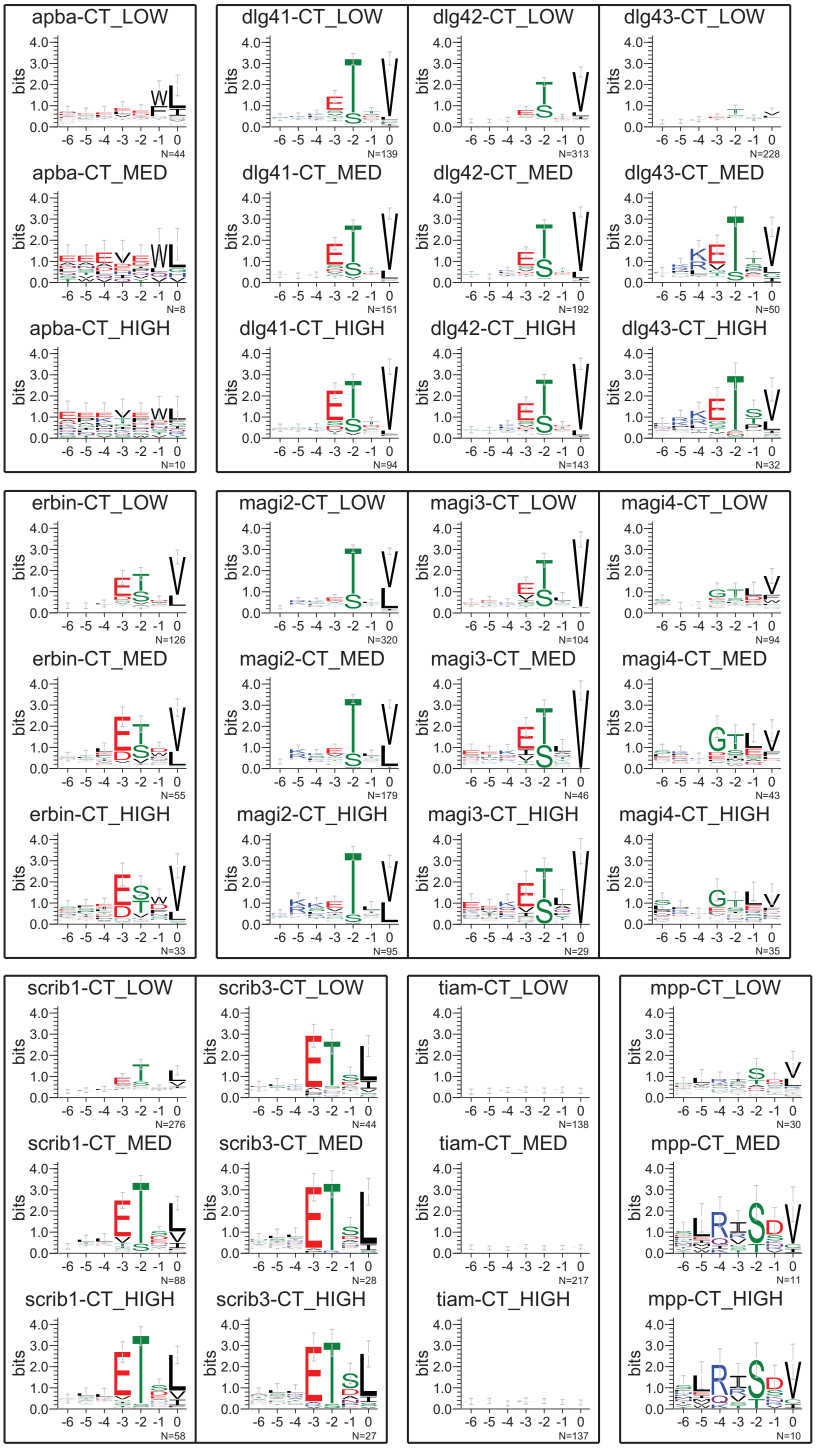

**Figure S3. PDZ specificities determined by MR-B1H selection and GFP sort using a 7aa Natural library.** Motifs represent peptides selected from the Natural library that consists of ~50,000 C-termini of human ORFs and splice variants[1]. These selected peptides were recovered from populations at various GFP levels. The name of the PDZ domain tested is labeled above each motif as well as the GFP level it represents (LOW, MED(medium), or HIGH).

A.

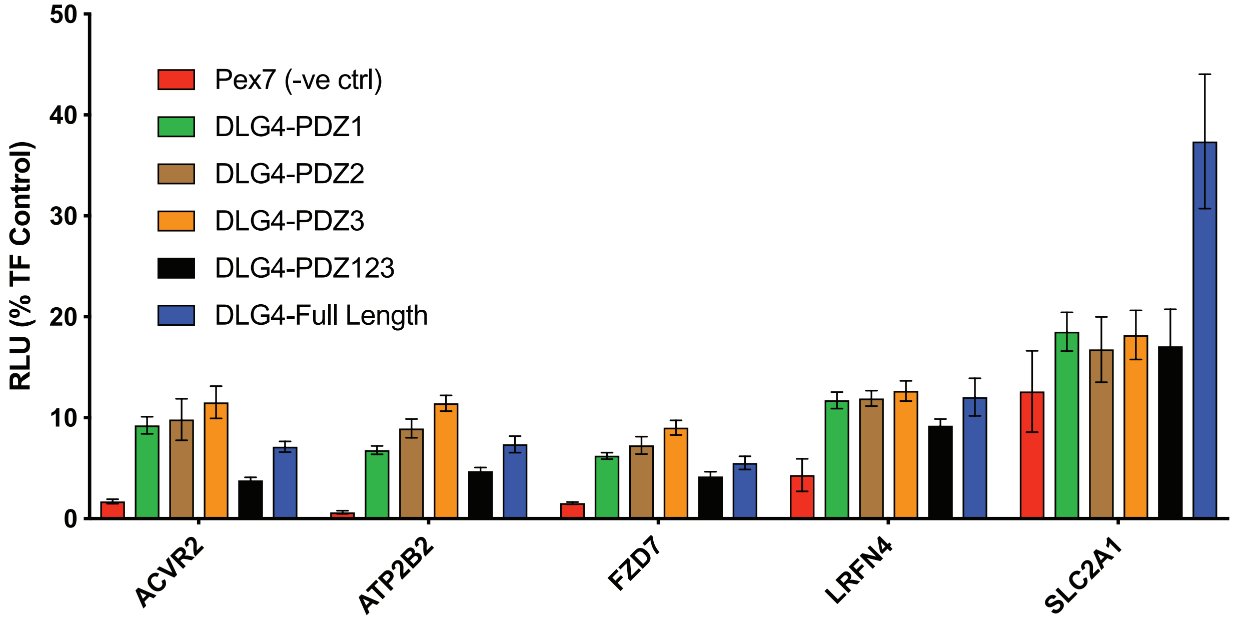

B.

| | DLG4-PDZ1 | DLG4-PDZ2 | DLG4-PDZ3 | DLG4-PDZ123 | DLG4-Full-Length |
|---|---|---|---|---|---|
| ACVR2B | +++ | +++ | +++ | - | +++ |
| ATP2B2 | +++ | +++ | +++ | + | +++ |
| FZD7 | + | +++ | +++ | - | + |
| LRFN4 | +++ | +++ | +++ | + | +++ |
| SLC2A1 | +++ | + | ++ | + | +++ |

**Figure S4. Validation of full-length PDZ interactions by MaMTH assay.** Full-length PSD-95 (DLG4) and truncated versions that include each or all of the PDZ domains were tested for their ability to bind full-length versions of predicted binding partners. **A**) The control construct (Pex7) or the PSD-95 variants were paired with the predicted full-length partners noted on the X-axis. For each pairing luciferase expression was measured as Relative Light Units (RLU) and normalized as the fraction of the output produced by a control transcription factor. **B**) A chart of *p*-values obtained from a 2way ANOVA (multiple comparisons) between the indicated interaction and the corresponding Pex7 negative control ($p \geq 0.05$ (-); $p<0.05$ (+); $p<0.005$ (++); $p<0.0005$ (+++)).

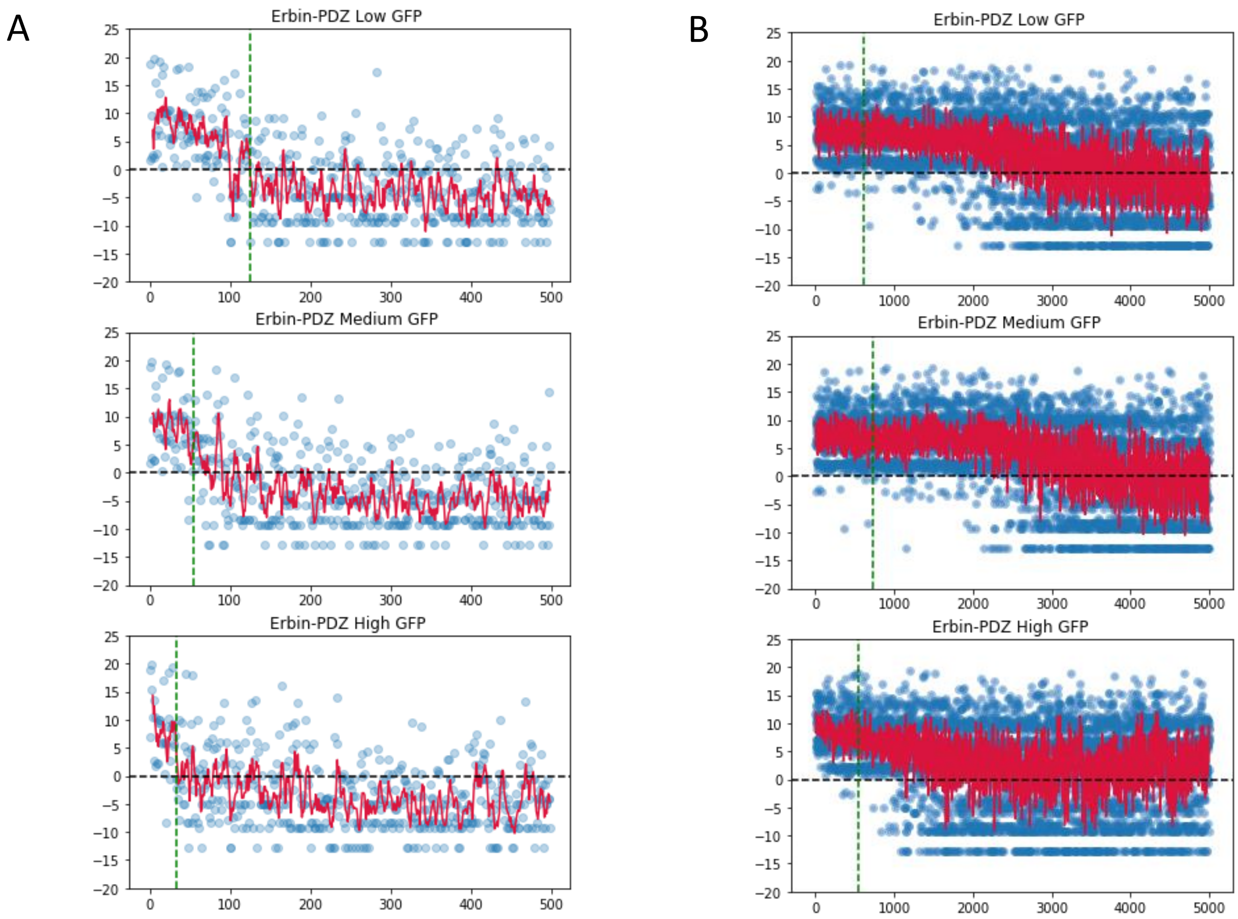

**Figure S5. Peptides ranked by read number (X-axis) and scored using a PWM built from a set of confirmed peptides (Y-axis) for the Erbin PDZ domain. A**) Peptide scores from the Natural library (cut-off = 3.4 normalized read counts) are shown. **B**) Peptide scores from the Random library (cut-off = top 5%) are shown. The dashed green line marks the cut-off threshold, and the red line is the moving average for the PWM score.

A.

pB1H2 zif268-Random library
CCATGGGTGCGGCCGCGGACTACAAGGATGACGACGACAAGTTCCGGACGGGCTCCAAGAC
ACCGCCTCACGGTACCGAGCGCCCATATGCTTGCCCTGTCGAGTCCTGCGATCGCCGCTTTT
CTCGCTCGGATGAGCTTACCCGCCATATCCGCATCCATACCGGTCAGAAGCCCTTCCAGTGTC
GAATCTGCATGCGTAACTTCAGTCGTAGTGACCACCTTACCACCCACATCCGCACCCACACAG
GCGAGAAGCCTTTTGCCTGTGACATTTGTGGGAGGAAGTTTGCCAGGAGTGATGAACGCAAG
AGGCATACCAAAATCCATCTGCGCGGATCCGGCGGTGGGAGCGGAGGCNNSNNSNNSNNSN
NSNNSNNSTAATAGGTCTAGA

Restriction sites (NcoI (encodes start codon) and XbaI)
Zif268 zinc fingers
Linker
Library

B.

pB1H2 zif268-Natural library
CCATGGGTGCGGCCGCGGACTACAAGGATGACGACGACAAGTTCCGGACGGGCTCCAAGAC
ACCGCCTCACGGAACCGAGCGCCCATATGCTTGCCCTGTCGAGTCCTGCGATCGCCGCTTTT
CTCGCTCGGATGAGCTTACCCGCCATATCCGCATCCATACCGGTCAGAAGCCCTTCCAGTGTC
GAATCTGCATGCGTAACTTCAGTCGTAGTGACCACCTTACCACCCACATCCGCACCCACACAG
GCGAGAAGCCTTTTGCCTGTGACATTTGTGGGAGGAAGTTTGCCAGGAGTGATGAACGCAAG
AGGCATACCAAAATCCATCTGCGCGGTACCGGCGCCCCGGTGGCGGANNNNNNNNNNNNN
NNNNNNNNTGATAAACCGATACAATAAAGGCTCTAGA

Restriction sites (KpnI and XbaI)
Zif268 zinc fingers
Linker
Library

C.

pGHUC ω-Erbin
ATGGCACGCGTAACTGTTCAGGACGCTGTAGAGAAAATTGGTAACCGTTTTGACCTGGTACTG
GTCGCCGCGCGTCGCGCTCGTCAGATGCAGGTAGGCGGAAAGGACCCGCTCGTACCGGAAG
AAAACGATAAAACCACTGTAATCGCGCTGCGCGAAATCGAAGAAGGTCTGATCAACAACCAGA
TCCTCGACGTTCGCGAACGCCAGGAACAGCAAGAGCAGGAAGCCGCTGAATTACAAGCCGTT
ACCGCTATTGCTGAAGGTCGTGCGGCGGCGGACTACAAGGATGACGACGACAAGTTCCGGAC
AGGCTCCAAGACACCTCCGCATTCTAGACGATCGGGCAGCCATATGGGCCATGAACTGGCGA
AACAGGAAATTCGCGTGCGCGTGGAAAAAGATCCGGAACTGGGCTTTAGCATTAGCGGCGGC
GTGGGCGGTCGTGGCAACCCGTTTCGCCCGGATGATGATGGCATTTTTGTGACCCGCGTGCA
GCCGGAAGGCCCGGCGAGCAAACTGCTGCAGCCGGGCGATAAAATTATTCAGGCGAACGGC
TATAGCTTTATTAACATTGAACATGGCCAGGCGGTGAGCCTGCTGAAAACCTTTCAGAACACC
GTGGAACTGATTATTGTGCGCGAAGTGAGCAGCTAATAGCTCGAG

Omega subunit
Linker
Restriction sites (XbaI and XhoI)
Erbin PDZ

Figure S6. Coding scheme for proteins employed. **A**) Zif268-Random library sequence is shown where NNS represents the code for each codon. **B**) Zif268-Natural library sequence is shown where NNN purely represents diversity but as these ~50,000 are natural peptides, NNN is NOT meant to indicate every possible combination. **C**) ω-Erbin sequence is shown.

### Methods

#### Library and plasmid construction

The Random library was constructed as previously described[3,4]. In short, 96 separate 15-cycle PCR reactions were performed using Zif268 as a template. The 3' oligonucleotide includes the code for the linker and random peptide region, stop codons and an XbaI site. In addition, the randomization scheme was requested to be hand mixed by IDT as our prior work had shown this approach led to a more true randomization (unpublished data). The low cycle number reduces the amplification of polymerase error and separate reactions minimizes the influence that any bias from a single reaction will have on the entire population when 96 reactions are pooled. The forward primer contained the NcoI restriction site that includes the start codon followed by an N-terminal flag tag. The pooled PCR product was digested with NcoI and XbaI and cloned into our expression vector that utilizes a LacUV5 promoter for expression.

The Natural library was constructed similarly, except the amplification only included the peptide region. Oligonucleotides were designed to amplify around the peptide region on the phagmid library previously investigated[1]. On the end of this sequence were placed cloning sites KpnI and XbaI.

Omega-PDZ fusions were created by restriction cloning into the pGHUC[6] vector where the omega-PDZ constructs are expressed from a lacUV5 promoter. PDZ domains can easily be cloned between XbaI and XhoI site to result in the omega fusions (**Figure S6**). PDZ domains were purchased commercially as dsDNA G-blocks from IDT.

**All base plasmids and complete sequences will be made available at addgene.*

#### Selections on plates

One μg of prey (zif268-peptide library fusion) was transformed with 1μg of concentrated bait (ω-PDZ fusion) reporter into the selection strain (*E. coli* – strain genotype: US0 ΔrpoZ, ΔhisB, ΔpyrF)[7]. Cells were recovered at 37°C first in 10ml SOC for 1 hour, and then in complete NM media for another hour, before washing with NM media that lacks histidine and concentration to 1ml for overnight storage. The washed cells are titrated by 10-fold serial dilutions on rich media plates supplemented with appropriate antibiotics to determine total number of transformants that took up both bait and prey plasmids (Kan and Amp, respectively). The following day, 5e7 cells were plated on 150mm x 15mm NM media plates (-histidine) containing 2, 5, or 10mM 3 Amino-1,2,4-triazole (3AT) and grown for 36 to 48 hours at 37°C. 5e7 cells are used to avoid crowding on the plates and we have not found that this severely limits the number of colonies that survive. However, if a PDZ domain were very specific it may require the investigation of larger fractions of the library to recover a substantial number of variants it is able to interact with. In this case we would plate 1e9 on large 245mm x 245mm square plates as we have done for other applications including the prep selections for sorting below (1e9 plated on 1mM 3AT for each PDZ to be sorted). Colony PCR was performed on surviving colonies and the sequences were quality checked and compiled into sequence logos using the WebLogo tool (http://weblogo.berkeley.edu/logo.cgi.

**Selection in liquid culture**

For liquid selection and sorting, ~1e9 cells were plated on large plates above at low stringency as detailed above. The surviving colonies per pooled and DNA harvested. This "round 1 DNA" was retransformed with the PDZ reporter under investigation, and treated just as a plate selection through the titration step. At this point, when transformants can be counted, $10^6$ cells are inoculated into 100mL NM media (-histidine) with 1mM 3AT and grown in shaking incubators at 37°C. We have found that the most informative data can be found just as the GFP expression approaches its maximum. However, it can be challenging to determine when that time point will be. Therefore, for each PDZ ~1mL of culture was spun down and resuspended in cold PBS + 0.5% FBS every 3 hours starting 12 hours after inoculation and ending 48 hours after inoculation. Samples were run on a SONY SH800 cell cytometer and the mean GFP was recorded for each time point. Populations of GFP positive cells (Low, medium, and high) just prior to saturation were recovered as well as a population of cells that were not expressing GFP (negatives). 1e6 cells from the four sub-populations of GFP were sorted: negative, low, medium, and high, with the mean GFP of each population ~5 fold higher than the preceding one. As noted, the time-point just prior to GFP saturation was chosen for sorting. ~5e5 cells/GFP population were plated on 2XYT agar with antibiotics and grown to single colony resolution (~10hours), whereupon cells were pooled, and libraries were extracted, barcoded, and sent for next-generation sequencing (NGS). In parallel, ~24 individual colonies were Sanger sequenced to compare to the NGS data.

**Next-generation Sequencing data processing**

After demultiplexing, all reads with an average quality Phred score of > 30 and mapping to the C-terminal library were selected and frequencies were calculated. Random library peptides represented by multiple nucleotide sequences were selected to distinguish peptides selected at the protein level (codon diversity but the same peptide) from background, breakthrough colonies (these would be a single DNA sequence as their breakthrough would have nothing to do with peptide enrichment). Reads not matching the selection criteria were marked as undetermined and discarded. Finally, the reads were then normalized to the total number of peptides read for that GFP population, and a scalar factor was applied for readability purposes.

To select the set of peptides that interact with the PDZ domains of the Natural library screenings, we required a minimum threshold of the normalized read count. In order to assign a cut-off value that provides the best separation between specific and non-specific values, we compared the peptides' ranks based on reads to PWM scoring derived from a set of biophysical validated peptides for each PDZ domain (**Figure S5a**). A consensus threshold of 3.4 normalized reads was applied to all the samples in order to maximize recall. The selected peptides include the majority of peptides with a high score in the PWM and the majority of the total reads for the sample. This model only seems to miss the optimal separation boundary in the low GFP level of Scribble PDZ1.

The same approach does not wholly apply to the random library (**Figure S5b**), mainly because of the size and diversity of the data collected. Consequently, it is challenging to find a consensus and simple threshold that generalizes for all the PDZ domains and samples. Here, to select the set of peptides from the random library that interact with the PDZ, we instead selected the top 5% of the peptides. This small group of peptides captures the majority of the reads (ranging from 65% to 85% percent of the total). This bimodal distribution of reads seems to be sufficient to provide a good split between non-specific binders and specific binders for all assayed samples.

### MaMTH validation assays

Entry clones were prepared in vector pDONR223, and moved into either MaMTH C-tagging bait Destination vector A1345 (for ACVR2B, ATP2B2, FZD7, LRFN4, and SLC2A1) or MaMTH N-tagging prey Destination vector A1245 (for DLG4, PEX7 and all PDZ constructs) using the Gateway Cloning Technology (Thermo Fisher Scientific). MaMTH assays were carried out as previously described[8]. In brief, HEK293 MaMTH reporter cells were seeded in 96-well plates and grown in DMEM (containing 10%FBS and 1%Penicillin-Streptomycin) overnight at 37˚C/5% $CO_2$. Cells were co-transfected with bait and prey plasmid constructs using X-tremeGENE 9 (Roche) as per manufacturer instructions. Five hours after transfection, media containing transfection reagent was removed and replaced with fresh DMEM/10%FBS/1%PS containing 0.5 μg/mL tetracycline to induce bait expression. After 24 hours growth at 37˚C/5% $CO_2$ luciferase activity was measured by chemiluminescence using a Clariostar microplate reader (BMG Labtech).

1 Ivarsson, Y. *et al.* Large-scale interaction profiling of PDZ domains through proteomic peptide-phage display using human and viral phage peptidomes. *Proc Natl Acad Sci U S A* **111**, 2542-2547, doi:10.1073/pnas.1312296111 (2014).

2 Tonikian, R. *et al.* A specificity map for the PDZ domain family. *PLoS Biol* **6**, e239, doi:10.1371/journal.pbio.0060239 (2008).

3 Persikov, A. V., Rowland, E. F., Oakes, B. L., Singh, M. & Noyes, M. B. Deep sequencing of large library selections allows computational discovery of diverse sets of zinc fingers that bind common targets. *Nucleic Acids Res* **42**, 1497-1508, doi:10.1093/nar/gkt1034 (2014).

4 Persikov, A. V. *et al.* A systematic survey of the Cys2His2 zinc finger DNA-binding landscape. *Nucleic Acids Res* **43**, 1965-1984, doi:10.1093/nar/gku1395 (2015).

5 Calderone, A., Castagnoli, L. & Cesareni, G. mentha: a resource for browsing integrated protein-interaction networks. *Nat Methods* **10**, 690-691, doi:10.1038/nmeth.2561 (2013).

6 Oakes, B. L. *et al.* Multi-reporter selection for the design of active and more specific zinc-finger nucleases for genome editing. *Nat Commun* **7**, 10194, doi:10.1038/ncomms10194 (2016).

7 Noyes, M. B. *et al.* A systematic characterization of factors that regulate Drosophila segmentation via a bacterial one-hybrid system. *Nucleic Acids Res* **36**, 2547-2560, doi:10.1093/nar/gkn048 (2008).

8 Petschnigg, J. *et al.* The mammalian-membrane two-hybrid assay (MaMTH) for probing membrane-protein interactions in human cells. *Nat Methods* **11**, 585-592, doi:10.1038/nmeth.2895 (2014).
